# AI-driven Classification of Heart Failure Preserved and Reduced Ejection Fraction Patients Using the Total Protein Approach

**DOI:** 10.1101/2024.11.25.625184

**Authors:** Alessio Lenzi, Luis B. Carvalho, Pedro A.D. Teigas-Campos, Denise Biagini, Silvia Ghimenti, Silvia Armenia, Nicola R. Pugliese, Stefano Masi, Fabio Di Francesco, Tommaso Lomonaco, Carlos Lodeiro, Jose-Luis Capelo-Martinez, Hugo M. Santos

**Author notes:** Authors contributed equally (Alessio Lenzi, Luis B. Carvalho).

## Abstract

Heart failure (HF) presents two major subtypes: HF with preserved ejection fraction (HFpEF) and HF with reduced ejection fraction (HFrEF), each one with distinct metabolic characteristics. This study utilized artificial intelligence, high-resolution mass spectrometry and the Total Protein Approach (TPA) to identify key features differentiating these subtypes. Aldolase A (ALDOA), a glycolytic enzyme, was found upregulated in HFrEF patients, reflecting an increased glycolysis pathway, while Arginase 1 (ARG1), a key enzyme in the urea cycle, was also elevated, indicating an increased urea pathway. In contrast, HFpEF patients showed TPA ALDOA and ARG1 levels similar to healthy controls. The combined use of ALDOA and ARG1 TPA values successfully classified 82% of patients (14 out of 17). Additionally, most HFpEF patients were over 80 years old, suggesting an age-related metabolic shift. The combination of ALDOA and ARG1 are promising biomarkers for distinguishing HFpEF and HFrEF using the TPA approach, with potential implications for targeted therapies.

## 1. Introduction

Heart failure (HF) is a complex clinical syndrome characterized by the heart’s inability to pump sufficient blood to meet the body’s needs. It encompasses a spectrum of diseases, including heart failure with reduced ejection fraction (HFrEF) and heart failure with preserved ejection fraction (HFpEF). While sharing common symptoms, these two subtypes diverge significantly in their pathophysiology, diagnostic criteria, and treatment strategies[1,2]. HFrEF, also known as systolic heart failure, is defined by an ejection fraction (EF) of less than 40%. This condition is primarily characterized by the heart’s reduced ability to contract and pump blood efficiently, often due to myocardial infarction or dilated cardiomyopathy [1]. Conversely, HFpEF, or diastolic heart failure, occurs when the heart contracts typically but exhibits impaired relaxation and filling during diastole, maintaining an EF of 50% or greater. Common etiologies for HFpEF include hypertension, obesity, and diabetes [3].

The diagnostic challenge in HF, particularly in HFpEF, is substantial. Accurate diagnosis requires differentiation between the two forms of HF, which often present with overlapping symptoms such as dyspnea, fatigue, and fluid retention. Traditional diagnostic tools, including echocardiography and natriuretic peptide levels, may not always provide clear distinctions. This diagnostic ambiguity can lead to misclassification and inappropriate management of the condition [4]. Moreover, the lack of specific biomarkers exacerbates the difficulty in diagnosing HF accurately. While biomarkers like B-type natriuretic peptide (BNP) and N-terminal pro-BNP (NT-proBNP) are elevated in HF, they do not effectively differentiate between HFrEF and HFpEF. Recent studies have explored other potential plasma biomarkers, such as galectin-3, ST2, and growth differentiation factor-15 (GDF-15), yet these markers still lack the specificity and sensitivity needed for routine clinical use [5,6].

Several urine biomarkers have been investigated for their potential role in diagnosing and managing heart failure. These biomarkers can provide valuable insights into heart failure patients’ physiological and pathological processes. Notable urine biomarkers for heart failure include Neutrophil Gelatinase-Associated Lipocalin (NGAL), which is associated with kidney injury and inflammation. Elevated levels of NGAL in urine have been linked to acute kidney injury in heart failure patients, indicating worsening renal function, a common comorbidity in heart failure. However, the specificity of NGAL for heart failure as opposed to other conditions causing kidney injury remains a challenge[7] [xx]. Natriuretic Peptides (BNP and NT-proBNP) are more commonly measured in blood but can also be detected in urine and serve as markers for heart failure severity and prognosis. However, their effectiveness in urine is less established than in blood measurements[8]. Albumin, specifically microalbuminuria, is an indicator of endothelial dysfunction and has been associated with an increased risk of heart failure. However, it can also be influenced by various other renal and systemic conditions, reducing its specificity for heart failure [9]. Kidney Injury Molecule-1 (KIM-1) levels in urine predict adverse outcomes in heart failure patients, reflecting kidney injury. Despite its potential, variations in KIM-1 levels due to non-cardiac renal injuries present diagnostic challenges [10]. N-Acetyl-β-D-glucosaminidase (NAG) is a marker of tubular damage in the kidneys. Elevated urinary NAG levels are associated with heart failure severity and renal impairment, but distinguishing its increase due to cardiac versus non-cardiac causes remains problematic. Liver-Type Fatty Acid-Binding Protein (L-FABP) levels in urine indicate kidney damage and have been linked to adverse cardiovascular outcomes in heart failure patients. Its clinical utility is still under investigation, as it also increases in various renal pathologies [11].

In this work, we have used analytical proteomics via high-resolution mass spectrometry and artificial intelligence to unveil biomarkers present in the urinary proteome of two groups of HF diagnosed as HFrEF and HFpEF, respectively. A healthy group was used as control. Results show that the complete proteome can be used to classify HFrEF and HFpEF and that AI can be used to speed data mining to find biomarkers straightforwardly, in this case, ALDOA Protein.

## 2. Methods

### 2.1. Patients

Urine samples were collected from three distinct patient cohorts: patients diagnosed with heart failure (HF) with reduced ejection fraction (HFrEF), patients with heart failure with preserved ejection fraction (HFpEF), and control group patients with no known cardiovascular conditions. Supplementary Material 1 contains patient-specific clinical and medical information.

All participants were told about the study and gave written consent before sample collection. The Ethics Committee of the University of Pisa accepted the study and the patient sample collection recruitment was carried out under the accepted ethical guidelines.

### 2.2. Sample Collection

Urine samples were collected in 100 mL containers, each containing 38 mg of boric acid (Sigma–Aldrich) to inhibit bacterial growth [12]. Samples exhibiting visible hematuria were excluded from the study and discarded. Following collection, the urine samples were centrifuged at 5000 × g for 20 minutes to eliminate cellular debris. The resulting supernatants were divided into 10 mL aliquots and stored at −60 °C until further analysis. After the completion of patient recruitment and sample collection, these supernatants were further aliquoted into smaller 1 mL portions and transported on dry ice for subsequent analyses.

### 2.3. Proteome Digestion

A modified filter-aided sample preparation (FASP) approach was used to digest different amounts of protein, ranging from 25 to 50 μg, equivalent to the volume of urine processed through a 10 kDa molecular weight cutoff (MWCO) membrane[13–15].

The protein retained in the membrane was washed with 200 μL of an 8 M urea 0.1 M Tris-HCl pH 8 (Solution A) solution, followed by centrifugation at 10,000 × g for 20 minutes. Protein reduction was performed by adding 200 μL of 50 mM dithiothreitol (DTT) in Solution A and incubating at 37 °C for 60 minutes, after which a centrifugation step was repeated for 20 minutes at 10,000 × g. The samples were then alkylated in the dark for 45 minutes by adding 100 μL of 50 mM iodoacetamide in Solution A, followed by centrifugation at 10,000 × g for 20 minutes. Samples were then washed twice with 0.07 M tetraethylammonium bromide (TEAB) and centrifuged at 10,000 × g for 20 minutes to ensure the solution added on top of the membrane flowed through it.

Protein digestion was performed overnight (~14 hours) at 37°C using 100 μL of trypsin solution at a 1:30 ratio in 0.07 M TEAB. The peptides were recovered by centrifugation at 10,000 × g for 20 minutes. The membrane was then washed twice with 100 μL of 3% (v/v) acetonitrile (ACN) containing 0.1% (v/v) formic acid (FA), followed by centrifugation at 10,000 × g for 20 minutes. The peptides were transferred to a 500-μL microcentrifuge tube, dried with a SpeedVac vacuum concentrator, and kept at −20°C for later analysis.

### 2.4. LC-MS/MS

Nano-liquid chromatography-tandem mass spectrometry (Nano-LC–MS/MS) was conducted utilizing an Ultimate 3000 ultra-high-performance nano-LC system (Thermo Scientific) in conjunction with an Ultra High-Resolution Quadrupole Time-of-Flight (UHR-QTOF) Impact HD mass spectrometer (Bruker Daltonics) equipped with a CaptiveSpray nanoBooster. A 3 μL peptide solution, containing 500 ng of peptides, was loaded onto a μPACTM Trapping column and desalted for 2.7 minutes using 2% (v/v) acetonitrile (ACN) in 0.1% formic acid (FA) at a flow rate of 15 μL/min. Chromatographic separation was conducted at 35 °C on an analytical column (200 cm μPACTM, PharmaFluidics) using a linear gradient at a flow rate of 500 nL/min. The mobile phase A (MPA) consisted of 0.1% (v/v) FA, and mobile phase B (MPB) was composed of 99.9% (v/v) ACN and 0.1% (v/v) FA. The gradient was as follows: from 0–2 min, 3% to 5% MPB (97% to 95% MPA); from 2–90 min, 5% to 35% MPB (95% to 65% MPA); from 90–100 min, 35% to 85% MPB (65% to 15% MPA); and from 100–120 min, 85% MPB (15% MPA). Mass spectrometry (MS) acquisition was configured for MS at 2 Hz, followed by MS/MS at 8–32 Hz, with a cycle time of 3.0 seconds. Active exclusion was applied, with exclusion after one spectrum and release after 2 minutes. Precursors were reconsidered if their intensity was threefold higher than the previous intensity, with a fragmentation intensity threshold set at 2,500 counts.

The mass spectrometry proteomics data have been deposited to the ProteomeXchange Consortium [16] (http://proteomecentral.proteomexchange.org) via the PRIDE partner repository [17].

### 2.5. Bioinformatic Analysis

#### 2.5.1. Relative Label-Free Quantification and Data Processing

Relative label-free quantification (LFQ) was obtained using the precursor signal intensity method and delayed normalization (MaxLFQ) on MaxQuant software V2.0.3.0 [18,19]. All raw files were processed in a single run with default parameters, and the database searches were performed using the MaxQuant integrated peptide search engine, Andromeda, against the human UniProt UP000005640_9606 database (20,600 sequences; 11,395,157 residues, downloaded on April 27, 2021) [20]. Searches were configured with cysteine carbamidomethylation as a fixed modification and N-terminal acetylation and methionine oxidation as variable modifications. The false discovery rate (FDR) was set to 0.01 for protein and peptide levels, with a minimum length of seven amino acids for peptides. Enzyme specificity was set as C-terminal to arginine and lysine, with a maximum of two missed cleavages allowed. Protein group LFQ intensities were processed using Perseus (version 2.1.1.0) with default settings [21]. Briefly, proteins were filtered to remove proteins identified with only a single modified peptide “only identified by site,” reverse sequences, and potential contaminants. Data was then log2-transformed and filtered to have a coverage with >70% in at least one group (Ctrl, HFrEF, HFpEF). Missing values were inputted from a Gaussian distribution using the parameters, with an upshift of 0.5 and a downshift of 1.8. Additional data were Quantile normalized using the R package limma add-on for Perseus [22,23]. Total protein abundance (TPA) methodology as also employed in this study, whereby the raw protein intensity values were divided by the total protein intensity in a sample and multiplied by the molecular weight of the respective identified proteins [24,25]. Protein data were filtered to remove proteins “only identified by site” peptides, reverse sequences, and potential contaminants.

#### 2.5.2. Machine Learning-based Artifical inteligence Classification and Feature Selection

A Random Forest classifier was employed to classify the different groups using the two already processed datasets (LFQ and TPA). Part of the computational workflow was optimized using a machine learning-based AI algorithm, which enhanced the computational efficiency and also fine-tuned model parameters, increasing the accurate predictions. Leave-One-Out Cross-Validation (LOO-CV) was used as the primary classification strategy, ideally for a robust approach in small datasets that iteratively train the model. Model accuracy was assessed by comparing true and predicted labels across all LOO-CV iterations, providing a comprehensive evaluation of the classification performance. Additionally, the Random Forest algorithm was used to assess the protein’s feature importance by extracting the top 20 proteins that contributed most to group differentiation. These key protein features were further visualized through box plots, illustrating their expression levels across the groups.

## 3. Results and discussion

Samples were divided into the following groups: (i) Heart Failure with Preserved Ejection Fraction (HFpEF), (ii) Heart Failure with Reduced Ejection Fraction (HFrEF), and (iii) Control (Ctrl). The groups were organized based on their technical replicates. AThe data were grouped into pairs based on their conditions: pEF-rEF, Ctrl-rEF, and Ctrl-pEF. This procedure is presented in a comprehensive scheme in Fig. 1a. Bioinformatics data treatment procedure is presented in section 2.5.

**Figure 1.**
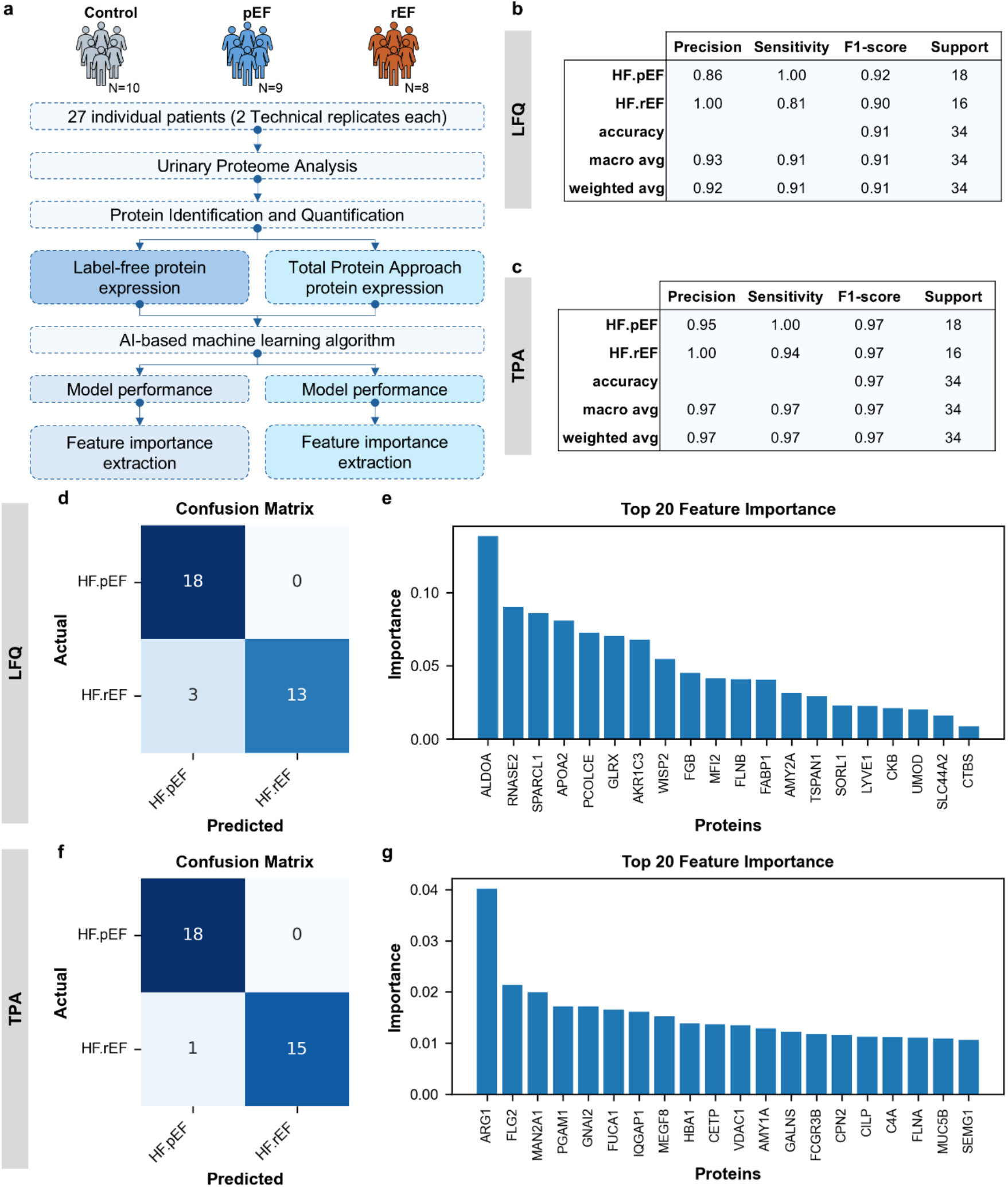
Heart failure patients’ classification. **a**. Workflow for data process and classification. **b**,**c**. Classification report using the different datasets, LFQ and TPA respectively. **d**,**f**. Confusion matrix for LFQ and TPA dataset, respectively, representing the classification performance of the Random Forest model classification model. Matrix illustrates the number of correct and incorrect predictions for each group. Rows correspond to actual group labels, and columns represent predicted labels. Darker shades indicate a higher number of correct classifications, while misclassifications are indicated in lighter shades. **e**,**g**. Bar plot depicting the top 20 most important features (proteins) for LFQ and TPA dataset, respectively, contributing to the Random Forest classification. Feature importance is ranked by the contribution to model accuracy, with higher bars indicating greater influence.

We compared conditions in pairs using the entire proteome, first based on protein expression levels and then using the Total Protein Approach (TPA). We chose the TPA method as the primary tool for classifying patients because it is faster for clinical purposes. Unlike LFQ, TPA allows individual patient classification without the need to analyze a full set of patients. The comparisons made were: Ctrl vs. HFpEF, Ctrl vs. HFrEF, and HFpEF vs. HFrEF. Using this approach, we analyzed the total proteomes of all patients to distinguish between the control, HFpEF, and HFrEF groups (Supplementary Material 2, Fig. 1, SM2 Fig.1). No statistically significant differences in protein levels were found between the groups.

Our main focus then shifted to distinguishing between patients with preserved (HFpEF) and reduced (HFrEF) ejection fractions, which is the key clinical challenge. As illustrated in Fig. 1b, using protein expression levels (LFQ) results in an overall classification accuracy of 91%, with a sensitivity of 100% for the HFpEF group and 81% for the HFrEF group. This is better visualized in Fig. 1d, where all 9 HFpEF patients and 7 out of 8 HFrEF patients were correctly classified. Out of the 17 patients, 2 were misclassified. The 20 most important proteins for classification are highlighted in Fig. 1e. Then we turned to make the classification but transforming the LFQ values into TPA values [24,25]. Results presented in Fig. 1c clearly show an improvement in the classification with an overall accuracy of 97% and a sensitivity of 100% and 94% for the HFpEF and the HFrEF groups, respectively. The confusion matrix presented in Fig. 1f shows that 9 out of 9 HFpEF patients and 7 out of 8 HFrEF patients are well-classified. Only 1 patient is confounded. The 20 proteins with the better classification power are shown in Fig, 1g. It is interesting to note that the top 20 classifier proteins identified using LFQ and TPA values differ.

### 3.1. Classifying heart failure using the Total Protein Approach

Using LFQ values, the best classifier protein identified was Aldolase A (ALDOA, Fig.1e), a key enzyme in glycolysis that catalyzes the conversion of fructose-1,6-bisphosphate into glyceraldehyde-3-phosphate and dihydroxyacetone phosphate. In heart failure, particularly in HFrEF, the heart undergoes significant metabolic changes, including a shift from fatty acid oxidation to a greater reliance on glycolysis for ATP production [26]. This shift underscores the importance of glycolytic enzymes like ALDOA, which play a crucial role in meeting the increased energy demands of the failing heart. The TPA values for ALDOA are presented in Fig. 3a. This adaptation is intended to boost glycolysis to compensate for the reduced efficiency of mitochondrial oxidative phosphorylation in low-oxygen conditions [26]. However, chronic reliance on glycolysis leads to increased lactate production and acidosis, which can worsen the progression of heart failure [27]. To ensure our data aligns with findings from the literature, we compared the glycolysis and gluconeogenesis pathways between the HFpEF and HFrEF groups. As shown in Fig. 2a, glycolysis was found to be downregulated in HFpEF patients, while it was upregulated in HFrEF patients. This result is consistent with previous studies [28]. While both HFpEF and HFrEF involve metabolic dysfunctions, HFrEF relies more heavily on glycolysis to meet energy demands, whereas HFpEF depends more on fatty acid metabolism and exhibits reduced glucose oxidation [29].

**Figure 2.**
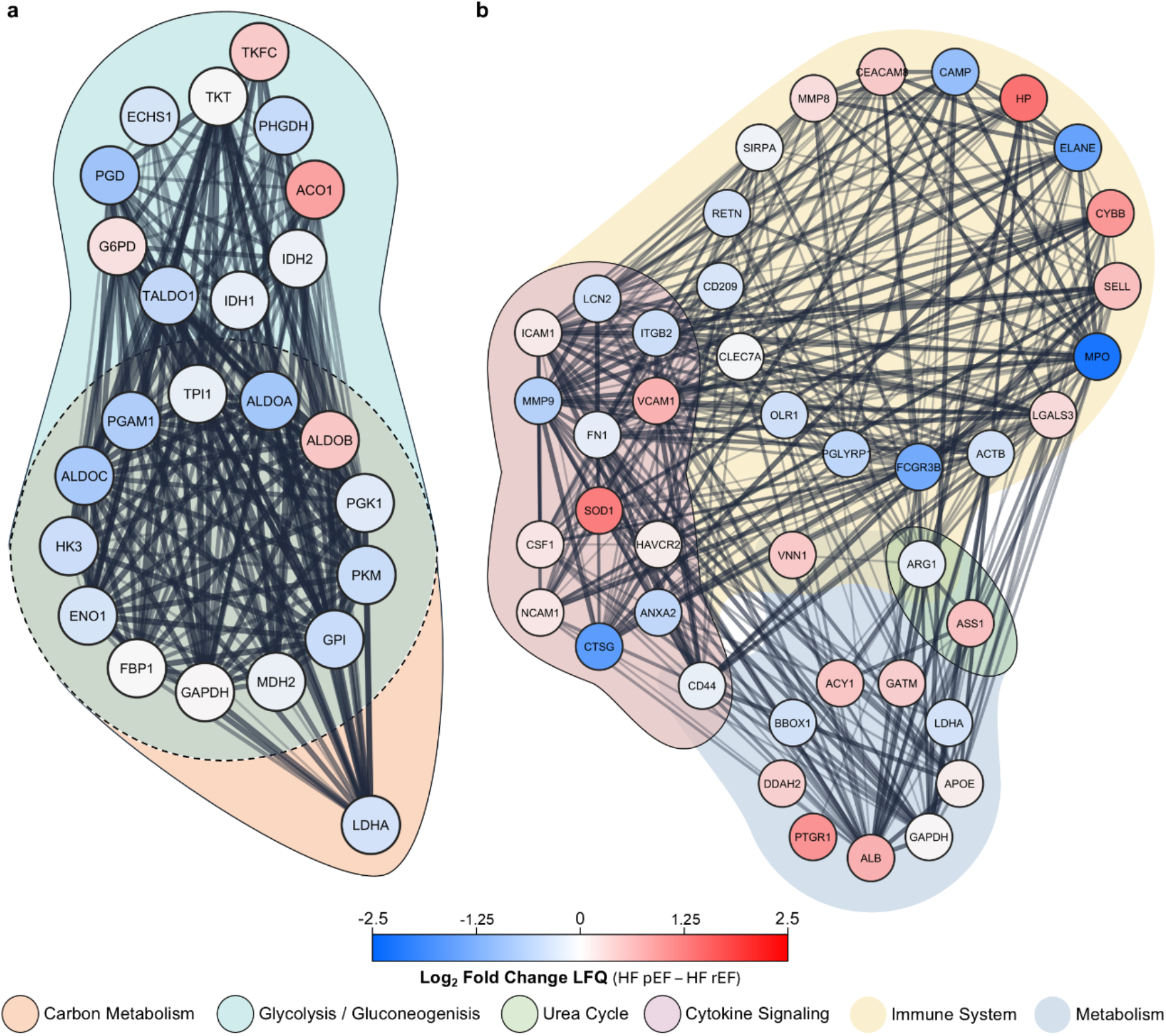
Protein-protein interaction pathway analysis of the target proteins ALDOA and ARG1. **a**. Protein network focusing of the proteins that directly connect with ALDOA and the enrichment pathways and biological processes that are directly correlated with the interactions. **b**. Protein network of the direct linked proteins to ARG1 and the enrichment pathways and processed directly link to the different connections.

**Figure 3.**
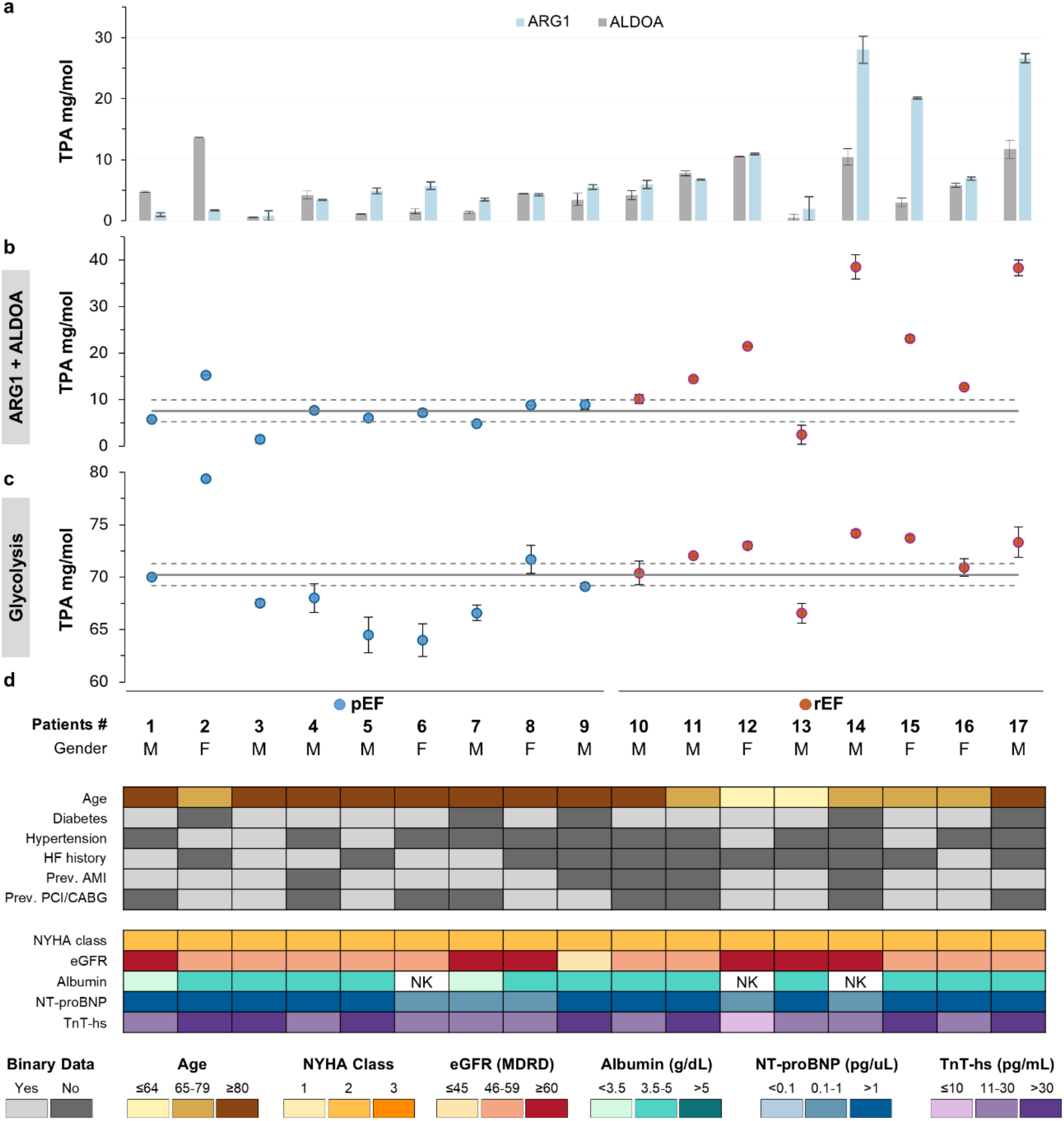
ALDOA and ARG1 expression levels integration into the different heart failure groups and medical data. **a**. TPA quantification expression levels of the ALDOA and ARG1 proteins. **b**. ALDOA and ARG combination expression levels to classify the HFpEF and HFrEF. **c**. Analysis of the combined proteins expression involved in the glycolysis process. **d**. Heat-map of the patient’s clinic information. Clinical parameters in heart failure: Age (50-89 years) reflects increasing risk with age. Binary data (Yes/No): Diabetes, hypertension, HF history, AMI, and PCI/CABG all elevate heart failure risk. NYHA Class (1-3) indicates symptom severity. eGFR (≤45 to ≥60) assesses kidney function. NT-proBNP (≤100 to >1000 pg/mL) and TnT-hs (≤10 to >30 pg/mL) reflect heart stress/injury. Albumin (<3.5 to >5.0 g/dL) indicates nutritional status and prognosis.

Since our findings on the glycolytic pathway (Fig. 2a) matched what has been reported in the literature, we selected proteins from this pathway that showed significant shifts (up or down) between the HFpEF and HFrEF groups, specifically Fructose-bisphosphate aldolase A (ALDOA), Fructose-bisphosphate aldolase C (ALDOC), Hexokinase-3 (HK3), and Phosphoglycerate mutase 1 (PGAM1). We then calculated the mean TPA value for these proteins using data from healthy individuals. Next, we calculated the same mean TPA value for each patient. The results, displayed in Fig. 3b, confirm a general trend: HFpEF patients’ TPA values are similar to those of the control group, while HFrEF patients tend to have higher TPA values.

When using the TPA values, Arginase 1 (ARG1) was recognized as the most effective classifier protein (Fig. 2b). This enzyme is essential in the urea cycle, facilitating the detoxification of ammonia produced during protein breakdown. Although primarily functioning in the liver, arginase is also expressed in various tissues, where it helps regulate arginine and nitric oxide levels. More recently, ARG1 has been detected in several blood cell types, including macrophages, endothelial cells, and smooth muscle cells. Studies have shown increased expression of ARG1 in cardiomyocytes [30]. Bekpinar et al. [31] proposed that ARG1 could serve as a biomarker, potentially increasing the risk of developing cardiac issues. In patients, elevated ARG1 levels are linked to a reduced left ventricular ejection fraction [32]. Consequently, arginase has emerged as a potential therapeutic target for treating heart failure, due to its role in worsening disease severity [32]. The TPA levels of ARG1 are presented in Fig. 3a.

Finally, we selected and combined the TPA values of the ALDOA and ARG1 proteins to differentiate between patients, as shown in Fig. 3c. For HFpEF patients, 8 out of 9 had values within or below the standard deviation of the average values for healthy individuals. For the HFrEF group, 6 out of 8 patients showed values higher than the standard deviation thresholds for healthy individuals, while one patient had values within the healthy range, and another had levels similar to the HFpEF group, making it indistinguishable.

Overall, from 17 patients, 6 out of 8 HFrEF patients and 8 out of 9 HFpEF patients were correctly classified, resulting in a total classification accuracy of 82% (14 out of 17 patients). Interestingly, 8 out of 9 HFpEF patients were over 80, as shown in Fig. 3d.

## 4. Conclusions

In this study, we identified Aldolase A (ALDOA) and Arginase 1 (ARG1) as the best TPA classifier proteins for distinguishing between HFpEF and HFrEF patients, both of which play critical roles in metabolic pathways relevant to heart failure. ALDOA, a key enzyme in glycolysis, was found to be upregulated in HFrEF, This finding is consistent with existing literature on metabolic shifts in heart failure. On the other hand, ARG1, primarily involved in the urea cycle, was also upregulated in HFrEF, indicating enhanced ammonia detoxification, a process tied to the breakdown of amino acids due to muscle wasting in heart failure.

Our data, supported by TPA value comparisons between healthy individuals and heart failure patients, confirmed significant metabolic differences between HFpEF and HFrEF groups. HFpEF patients generally exhibited the glycolysis pathway closer to healthy controls, while HFrEF patients showed elevated levels, particularly for proteins involved in glycolysis and the urea cycle. This metabolic divergence was further reinforced when combining the TPA values of ALDOA and ARG1, which successfully classified 82% of the patients. Additionally, an age-related trend was noted, with most HFpEF patients being older than 80 years, highlighting a potential link between age and metabolic regulation in heart failure. Overall, our findings suggest that ALDOA and ARG1 could serve as promising biomarkers for differentiating heart failure subtypes and potentially guiding targeted therapies for managing metabolic dysfunctions in heart failure patients.

## 6. Supplementary Data

Supplementary material 1: Detailed clinical patient data

Supplementary material 2: SM2 Fig1 (Proteome analysis of HFpEF, and HFrEF patients groups).

## 7. Author Contributions

### Contributions List

Conceptualization: Luis B. Carvalho; Jose-Luis Capelo-Martinez; Hugo M. Santos

Data curation: Luis B. Carvalho; Alessio Lenzi Experimental; Hugo M. Santos

Formal analysis: Luis B. Carvalho; Pedro A.D. Teigas-Campos; Hugo M. Santos

Funding acquisition: Carlos Lodeiro; Jose-Luis Capelo-Martinez; Hugo M. Santos

Investigation: Alessio Lenzi Experimental; Hugo M. Santos

Methodology: Alessio Lenzi Experimental; Luis B. Carvalho; Hugo M. Santos

Project administration: Jose-Luis Capelo-Martinez; Hugo M. Santos

Resources: Denise Biagini; Silvia Ghimenti; Silvia Armenia; Nicola R. Pugliese; Stefano Masi; Fabio Di

Francesco; Tommaso Lomonaco; Carlos Lodeiro; Jose-Luis Capelo-Martinez; Hugo M. Santos

Software: Luis B. Carvalho; Pedro A.D. Teigas-Campos

Supervision: Hugo M. Santos

Validation: Hugo M. Santos

Visualization: Luis B. Carvalho

Writing – original draft: Luis B. Carvalho; Jose-Luis Capelo-Martinez; Hugo M. Santos

Writing – review & editing: Alessio Lenzi Experimental; Luis B. Carvalho; Pedro A.D. Teigas-Campos; Denise Biagini; Silvia Ghimenti; Silvia Armenia; Nicola R. Pugliese; Stefano Masi; Fabio Di Francesco; Tommaso Lomonaco; Carlos Lodeiro; Jose-Luis Capelo-Martinez; Hugo M. Santos

## 8. Acknowledgments

PROTEOMASS Scientific Society is acknowledged by the funding provided to the Laboratory for Biological Mass Spectrometry Isabel Moura (#PM001/2019 and #PM003/ 2016). This work received support from Fundação para a Ciência e a Tecnologia and Ministério da Ciência, Tecnologia e Ensino Superior (FCT/MCTES) through the projects LA/P/0008/2020 DOI 10.54499/LA/P/0008/2020, UIDP/50006/2020 DOI 10.54499/UIDP/50006/2020 and UIDB/50006/2020 DOI 10.54499/UIDB/50006/2020. H.M.S. acknowledges the Associate Laboratory for Green Chemistry-LAQV (LA/P/0008/2020) DOI 10.54499/LA/P/0008/2020 funded by FCT/MCTES for his research contract. L.B.C. are supported by PROTEOMASS Scientific Society and the FCT/MCTES (2023.00528.BD).

## References

[1] P. Ponikowski, A.A. Voors, S.D. Anker, H. Bueno, J.G.F. Cleland, A.J.S. Coats, V. Falk, J.R. González-Juanatey, V.-P. Harjola, E.A. Jankowska, M. Jessup, C. Linde, P. Nihoyannopoulos, J.T. Parissis, B. Pieske, J.P. Riley, G.M.C. Rosano, L.M. Ruilope, F. Ruschitzka, F.H. Rutten, P. van der Meer, 2016 ESC Guidelines for the diagnosis and treatment of acute and chronic heart failure, Eur Heart J 37 (2016) 2129–2200. 10.1093/eurheartj/ehw128.

[2] S.J. Shah, D.H. Katz, S. Selvaraj, M.A. Burke, C.W. Yancy, M. Gheorghiade, R.O. Bonow, C.-C. Huang, R.C. Deo, Phenomapping for novel classification of heart failure with preserved ejection fraction, Circulation 131 (2015) 269–279. 10.1161/CIRCULATIONAHA.114.010637.

[3] S.M. Dunlay, V.L. Roger, Understanding the epidemic of heart failure: Past, present, and future, Curr Heart Fail Rep 11 (2014) 404–415. 10.1007/s11897-014-0220-x.

[4] W.J. Paulus, C. Tschöpe, A Novel paradigm for heart hailure with preserved ejection fraction, J Am Coll Cardiol 62 (2013) 263–271. 10.1016/j.jacc.2013.02.092.

[5] R.R. van Kimmenade, J.L. Januzzi, P.T. Ellinor, U.C. Sharma, J.A. Bakker, A.F. Low, A. Martinez, H.J. Crijns, C.A. MacRae, P.P. Menheere, Y.M. Pinto, Utility of amino-terminal pro-brain natriuretic peptide, galectin-3, and apelin for the evaluation of patients with acute heart failure, J Am Coll Cardiol 48 (2006) 1217–1224. 10.1016/j.jacc.2006.03.061.

[6] T.J. Wang, K.C. Wollert, M.G. Larson, E. Coglianese, E.L. McCabe, S. Cheng, J.E. Ho, M.G. Fradley, A. Ghorbani, V. Xanthakis, T. Kempf, E.J. Benjamin, D. Levy, R.S. Vasan, J.L. Januzzi, Prognostic utility of novel biomarkers of cardiovascular stress, Circulation 126 (2012) 1596–1604. 10.1161/CIRCULATIONAHA.112.129437.

[7] D. Bolignano, V. Donato, G. Coppolino, S. Campo, A. Buemi, A. Lacquaniti, M. Buemi, Neutrophil Gelatinase–Associated Lipocalin (NGAL) as a marker of kidney damage, American Journal of Kidney Diseases 52 (2008) 595–605. 10.1053/j.ajkd.2008.01.020.

[8] F.H. Epstein, E.R. Levin, D.G. Gardner, W.K. Samson, Natriuretic peptides, New England Journal of Medicine 339 (1998) 321–328. 10.1056/NEJM199807303390507.

[9] H.C. Gerstein, Albuminuria and risk of cardiovascular events, death, and heart failure in diabetic and nondiabetic individuals, JAMA 286 (2001) 421. 10.1001/jama.286.4.421.

[10] W.K. Han, V. Bailly, R. Abichandani, R. Thadhani, J. V. Bonventre, Kidney Injury Molecule-1 (KIM-1): A novel biomarker for human renal proximal tubule injury, Kidney Int 62 (2002) 237– 244. 10.1046/j.1523-1755.2002.00433.x.

[11] S.E. Nielsen, T. Sugaya, P. Hovind, T. Baba, H.-H. Parving, P. Rossing, Urinary liver-type fatty acid-binding protein predicts progression to nephropathy in type 1 diabetic patients, Diabetes Care 33 (2010) 1320–1324. 10.2337/dc09-2242.

[12] V. Thongboonkerd, P. Saetun, Bacterial overgrowth affects urinary proteome analysis: Recommendation for centrifugation, temperature, duration, and the use of preservatives during sample collection, J Proteome Res 6 (2007) 4179–4187. 10.1021/pr070311+.

[13] J.R. Wiśniewski, A. Zougman, N. Nagaraj, M. Mann, Universal sample preparation method for proteome analysis, Nat Methods 6 (2009) 359–362. 10.1038/nmeth.1322.

[14] J.R. Wiśniewski, Filter aided sample preparation – A tutorial, Anal Chim Acta 1090 (2019) 23–30. 10.1016/j.aca.2019.08.032.

[15] L.B. Carvalho, J.-L. Capelo-Martínez, C. Lodeiro, J.R. Wiśniewski, H.M. Santos, Ultrasonic-based filter aided sample preparation as the general method to sample preparation in proteomics, Anal Chem 92 (2020) 9164–9171. 10.1021/acs.analchem.0c01470.

[16] E.W. Deutsch, N. Bandeira, Y. Perez-Riverol, V. Sharma, J.J. Carver, L. Mendoza, D.J. Kundu, S. Wang, C. Bandla, S. Kamatchinathan, S. Hewapathirana, B.S. Pullman, J. Wertz, Z. Sun, S. Kawano, S. Okuda, Y. Watanabe, B. MacLean, M.J. MacCoss, Y. Zhu, Y. Ishihama, J.A. Vizcaíno, The ProteomeXchange consortium at 10 years: 2023 update, Nucleic Acids Res 51 (2023) D1539– D1548. 10.1093/nar/gkac1040.

[17] Y. Perez-Riverol, J. Bai, C. Bandla, D. García-Seisdedos, S. Hewapathirana, S. Kamatchinathan, D.J. Kundu, A. Prakash, A. Frericks-Zipper, M. Eisenacher, M. Walzer, S. Wang, A. Brazma, J.A. Vizcaíno, The PRIDE database resources in 2022: a hub for mass spectrometry-based proteomics evidences, Nucleic Acids Res 50 (2022) D543–D552. 10.1093/nar/gkab1038.

[18] J. Cox, M. Mann, MaxQuant enables high peptide identification rates, individualized p.p.b.-range mass accuracies and proteome-wide protein quantification, Nat Biotechnol 26 (2008) 1367–1372. 10.1038/nbt.1511.

[19] J. Cox, M.Y. Hein, C.A. Luber, I. Paron, N. Nagaraj, M. Mann, Accurate proteome-wide label-free quantification by celayed normalization and maximal peptide ratio extraction, termed MaxLFQ,Molecular & Cellular Proteomics 13 (2014) 2513–2526. 10.1074/mcp.M113.031591.

[20] J. Cox, N. Neuhauser, A. Michalski, R.A. Scheltema, J. V. Olsen, M. Mann, Andromeda: A peptide search engine integrated into the MaxQuant environment, J Proteome Res 10 (2011) 1794–1805. 10.1021/pr101065j.

[21] S. Tyanova, T. Temu, P. Sinitcyn, A. Carlson, M.Y. Hein, T. Geiger, M. Mann, J. Cox, The Perseus computational platform for comprehensive analysis of (prote)omics data, Nat Methods 13 (2016) 731–740. 10.1038/nmeth.3901.

[22] S. Yu, D. Ferretti, J.P. Schessner, J.D. Rudolph, G.H.H. Borner, J. Cox, Expanding the Perseus software for omics data analysis with custom plugins, Curr Protoc Bioinformatics 71 (2020). 10.1002/cpbi.105.

[23] M.E. Ritchie, B. Phipson, D. Wu, Y. Hu, C.W. Law, W. Shi, G.K. Smyth, Limma powers differential expression analyses for RNA-sequencing and microarray studies, Nucleic Acids Res 43 (2015) e47–e47. 10.1093/nar/gkv007.

[24] J.R. Wiśniewski, Label-free and standard-free absolute quantitative proteomics using the “total protein” and “proteomic ruler” approaches, in: 2017: pp. 49–60. 10.1016/bs.mie.2016.10.002.

[25] S. Jorge, J.L. Capelo, W. LaFramboise, S. Satturwar, D. Korentzelos, S. Bastacky, G. Quiroga-Garza, R. Dhir, J.R. Wiśniewski, C. Lodeiro, H.M. Santos, Absolute quantitative proteomics using the total protein approach to identify novel clinical immunohistochemical markers in renal neoplasms, BMC Med 19 (2021) 196. 10.1186/s12916-021-02071-9.

[26] G.L. Semenza, Hypoxia-inducible factors in physiology and medicine, Cell 148 (2012) 399–408. 10.1016/j.cell.2012.01.021.

[27] M. Bayeva, M. Gheorghiade, H. Ardehali, Mitochondria as a therapeutic target in heart failure, J Am Coll Cardiol 61 (2013) 599–610. 10.1016/j.jacc.2012.08.1021.

[28] N. Koleini, M. Meddeb, L. Zhao, M. Keykhaei, S. Kwon, F. Farshidfar, V.S. Hahn, E.L. Pearce, K. Sharma, D.A. Kass, Landscape of glycolytic metabolites and their regulating proteins in myocardium from human heart failure with preserved ejection fraction, Eur J Heart Fail (2024). 10.1002/ejhf.3389.

[29] P. Li, H. Zhao, J. Zhang, Y. Ning, Y. Tu, D. Xu, Q. Zeng, Similarities and differences between HFmrEF and HFpEF, Front Cardiovasc Med 8 (2021). 10.3389/fcvm.2021.678614.

[30] S.F.A. Shah, S. Akram, T. Iqbal, S. Nawaz, M.A. Rafiq, S. Hussain, Association analysis between ARG1 gene polymorphisms and idiopathic dilated cardiomyopathy, Medicine 98 (2019) e17694. 10.1097/MD.0000000000017694.

[31] S. Bekpinar, F. Gurdol, Y. Unlucerci, S. Develi, A. Yilmaz, Serum levels of arginase I are associated with left ventricular function after myocardial infarction, Clin Biochem 44 (2011) 1090– 1093. 10.1016/j.clinbiochem.2011.06.003.

[32] Z. Li, L. Wang, Y. Ren, Y. Huang, W. Liu, Z. Lv, L. Qian, Y. Yu, Y. Xiong, Arginase: shedding light on the mechanisms and opportunities in cardiovascular diseases, Cell Death Discov 8 (2022) 413. 10.1038/s41420-022-01200-4.

